# ClumPyCells resolves spatial aggregation in complex tissues overcoming size biases

**DOI:** 10.64898/2026.03.26.714529

**Authors:** Zetong Zhao, Jiabo Cui, Alicia G. Aguilar-Navarro, Maryam Monajemzadeh, Qing Chang, Ziyu Chen, Hubert Tsui, Eugenia Flores-Figueroa, Gregory W. Schwartz

## Abstract

The spatial arrangement of cells within a tissue microenvironment shapes their interactions and cell states, which are essential for tissue development, homeostasis, and disease. Spatial -omics technologies can precisely map the location of each cell within complex tissue structures, while also profiling their protein content and transcriptional diversity. Various approaches have been developed to analyze spatial patterns of cell aggregation, repulsion, or random distribution within tissues. However, differences in cell morphology within a tissue can introduce significant bias. Cell size in particular is not accounted for and introduces challenges when quantifying the aggregation of cells or their molecular features. To overcome such limitations, we present ClumPyCells: a statistical framework that measures cell and marker aggregation within tissue while correcting for size morphology. ClumPyCells enables interpretation of cell aggregation, bypassing interfering cell types or tissue regions unrelated to the desired spatial correlation. We demonstrate the capabilities of ClumPyCells across several tumor types, including melanoma and colorectal cancer, and spatial -omics technologies such as spatial transcriptomics and proteomics, while benchmarking how cell-size differences contribute to misinterpretations. By correcting for disruptive cell types within a region of interest, ClumPyCells will determine new tissue patterns and structures without morphological interference.

## Introduction

Although cells exhibit a degree of cell autonomous functionality through intrinsic signaling, together they self-organize into higher-order structures to coordinate complex tissue function. Through dense networks of cellular interaction and communication, cells aggregate to control tissue homeostasis through mechanisms such as inflammation, angiogenesis, and migration.^1–3^ Tumors alter these structures to create a new microenvironment that facilitates growth.^4^ To better understand such processes, technologies providing tissue imaging paired with molecular measurements report critical information on the spatial context of each individual cell in situ. Each imaging technology measures different -omic modalities of cells, with a focus on proteomic (e.g. imaging mass cytometry (IMC),^5^ cyclic immunofluorescence,^6^ codetection by indexing^7^) and transcriptomic (e.g. spatial transcriptomics such as Visium HD,^8^ Stereo-seq,^9^ NICHE-seq,^10^ and Slide-seq^11^) or both (e.g. Digital Spatial Profiling^12^) data modalities. While these advances will enable crucial new insights into spatial biology and disease, they introduce major challenges to data interpretability.

Although existing analytical techniques try to bridge the gap between spatial and molecular information generated from these sophisticated technologies, there are significant challenges that limit their capabilities. Many methods are tailored towards specific data modalities, such as gene expression only from spatial transcriptomic data.^13–15^ In the context of spatial transcriptomics, single-cell spatial information aids in the annotation of cellular composition and the identification of cellular interactions.^13^ Other methods, like Starfysh^16^ and Giotto,^14^ provide cell-niche detection with spatial transcriptomics and proteomics. Utilizing IMC data, ImaCytE visualizes cellular niches among different clusters, but does not directly spatially correlate cell types within tumor microenvironments.^17^ Methods quantifying cellular proximity generally use spatial statistics such as Squidpy,^18^ spatstat,^19^ and SPIAT.^20^ However, common among all of these algorithms is the lack of correction for cellular morphology, especially cell size.^21^ This oversight can lead to misleading interpretations — for example, adipocytes can be as large as 300 µm in diameter, dwarfing the size of cytotoxic T cells at 5 µm – 10 µm.^22,23^ Such large cells occupy a massive amount of space, leading spatial aggregation methods to assume that all other cells are aggregated together when they could be uniformly distributed with no significant aggregation. While current programs, including the popular CellProfiler,^24^ segment cells to identify their precise position in the tissue and report their overall cell size, the latter information is generally unused. This highlights the need for more comprehensive approaches that integrate both spatial and morphological features.

To overcome these limitations, we introduce ClumPyCells as a data-modality- and imaging-technology-agnostic technique to quantify cellular and molecular aggregation while also correcting for diverse cellular morphologies. Inspired by tree species aggregation measurements from ecology,^25^ we used a novel implementation of the mark correlation function for cell-based analysis.^26^ Using the size correction feature, ClumPyCells reduces the effect of large cells for a better representation of the underlying distribution of cells in tissue.

Through exhaustive benchmarking, we here demonstrate the effect of large cell disruption to expected spatial statistics. We compared ClumPyCells to other commonly-used methods, finding superior accuracy with our size-corrected technique. We validated ClumPyCells on melanoma tissue with different responses to immunotherapy. Using ClumPyCells on bone marrow tissue collected from a new cohort of acute myeloid leukemia (AML) patients, we identified key spatial factors classifying AML from normal bone marrow. To illustrate the flexibility of our method, we identified T-cell aggregation across colorectal cancer tissue measured with Visium HD. Built with compatibility in mind for Squidpy and Anndata, ClumPyCells is a powerful method for cell and feature aggregation analysis that exemplifies the need to account for cellular morphology in spatial biology. ClumPyCells is publicly available at https://github.com/schwartzlab-methods/ClumPyCells.

## Results

### ClumPyCells enables size-corrected aggregation measurements for cell types and features

Cell segmentation provides both spatial and morphological information within the tissue microenvironment. Existing spatial analysis tools fail to account for cell size, which can distort the understanding of cellular organization in tissues. Inspired by ecological techniques used to study tree aggregation, we introduce ClumPyCells, which corrects for these size-related biases. ClumPyCells is a Python package that applies mark correlation functions for cellular and molecular correlation analysis in tumor microenvironments (Figure 1). By adding a size correction feature, it is a powerful tool for better distribution representation. This approach enhances the accuracy of downstream analyses by improving the interpretation of cellular interactions and tissue organization.

**Figure 1:**
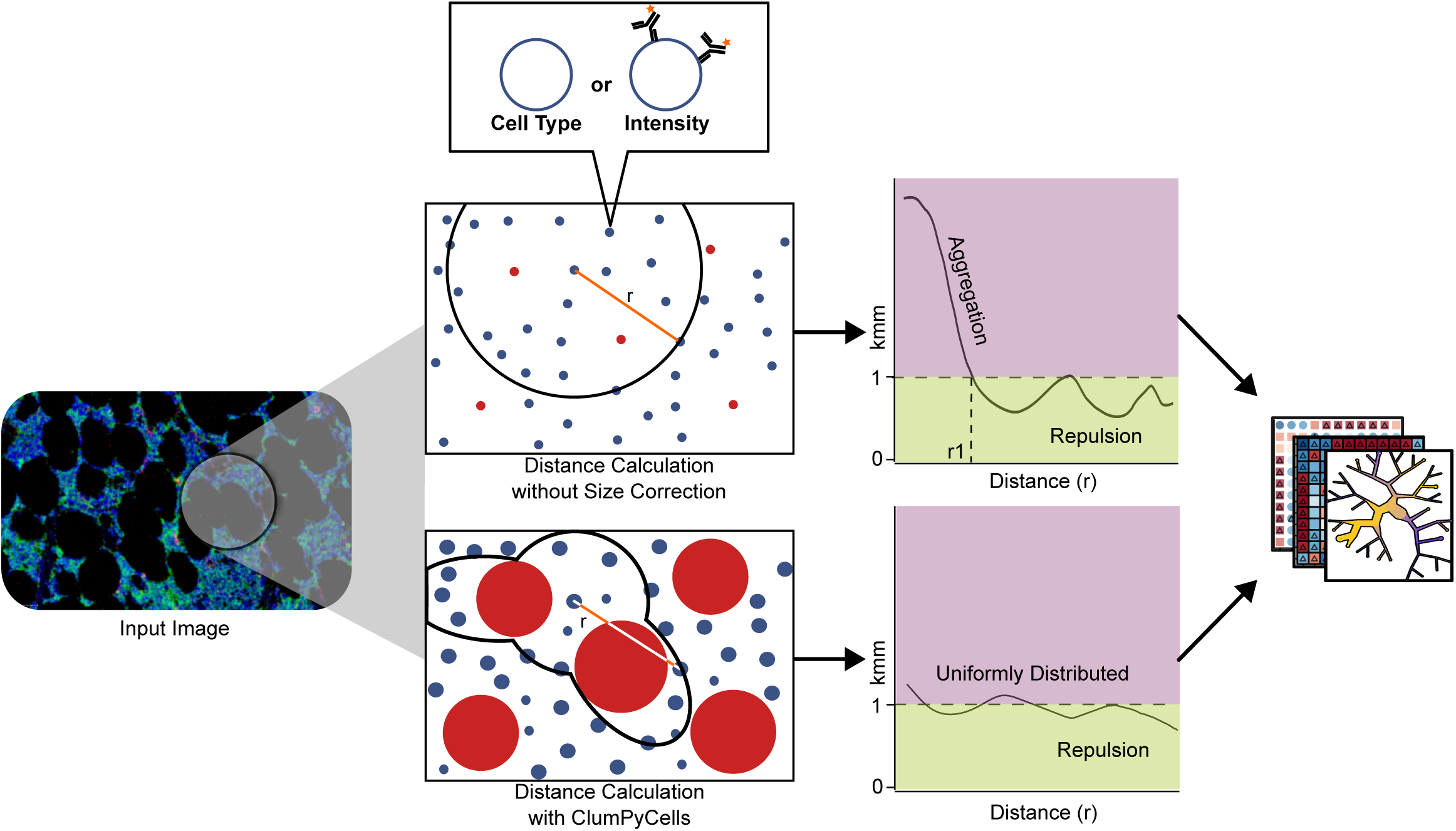
Overview of ClumPyCells. Cells from biological images are labeled by cell type or marker intensity. ClumPyCells supports two different analyses — with and without size correction. When large cells are present, ClumPyCells is able to give a distribution measurement of small cells without the disruptive effect of large cells. ClumPyCells outputs measurements correspond to different inter-cellular distances (*r*) which can be applied to further analysis.

ClumPyCells uses spatial information and labels of cells as input from technologies such as IMC and Visium HD. Spatial information includes Cartesian coordinates of cells, their diameters, and the window of analysis within the region of interest. Labels of the cells can be discrete labels like cell type and gene, or continuous labels like protein abundance or gene expression. ClumPyCells will output aggregation measurements for cells within the selected area.

ClumPyCells measures aggregation based on the mark correlation function, which uses inter-point distances within a population to calculate spatial correlation between labels such as cell type and protein abundance (“marks”). The method takes different distances depending on the labels as input and outputs correlation scores as a function of distance. Conceptually, the function compares the expected value of all the inter-point distances with a label by counting cell centers captured within a disc surrounding cells of the same label compared to the expected value of all inter-point distances between any label of the same-sized disc. However, this traditional mark correlation function is built on a point processing theory, which assumes each point (cell) is identical in size. This assumption does not hold in diseases such as cancer, where the tumor microenvironment contains cells with vastly different sizes which alter distances between cell membranes. More specifically, the inter-point distance between two cell centers does not represent the inter-cell distance in such an environment. ClumPyCells overcomes this limitation using a size-correction algorithm. We define a new inter-cell distance by taking the distance between two cell centers, and subtracting the radius of the two cells based on an estimated cell size from the center to the membrane. ClumPyCells may also remove cells or regions which could potentially distort the cell distribution in a tumor microenvironment by adjusting the inter-cell distance to exclude the portion passing through such regions. Importantly, ClumPyCells is a flexible model that supports AnnData format as input, and is fully compatible with Squidpy.

### Size-correction with ClumPyCells is robust to large cell interference

To assess the disruption effect when introducing large cells, we generated ground-truth synthetic data sets for comparative analysis with ClumPyCells (Figure 2). We generated 100 randomly-generated synthetic tissues comprised of three cell types, two small and one large (30 times larger than the small cells, mimicking adipocyte relative sizes), across four aggregation layouts to evaluate the spatial proximity between the two types of small cells (Figure 2a). We determined aggregation in one of two ways: one layout had only small cells located based on an intermixed uniform distribution, while the other layout grouped each cell type within specific regions of the tissue. Using these two layouts, we also injected large cells into the layout to measure their effect on aggregation.

**Figure 2:**
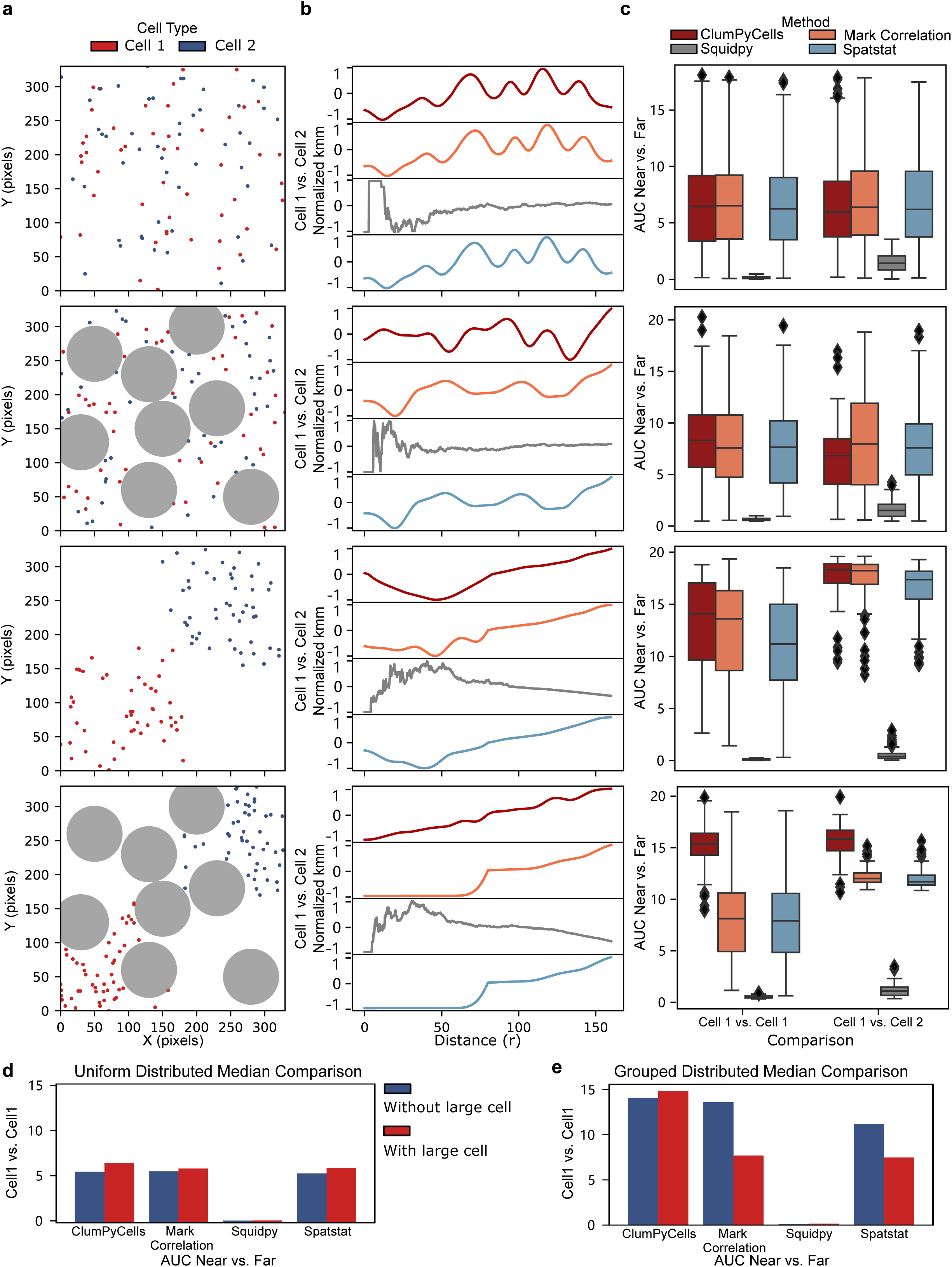
ClumPyCells is robust against large-cell presence in synthetic tissue benchmarks. **a**–**c**, Four synthetic experiment layouts of two small and one large cell type where each row is one of the four types of cell distributions (top to bottom: uniform, grouped, uniform with large cells, grouped with large cells). **a**, Representative synthetic experiments out of *n* 100 for each layout. **b**, Min-mid-max normalized output curves measuring aggregation between the two small cell types, where each row is a different algorithm at different distances (top to bottom: without size correction, ClumPyCells, Squidpy, and spatstat). **c**, Box-and-whisker plots (center line, median; box limits, upper (75^th^) and lower (25^th^) percentiles; whiskers, 1.5 interquartile range; points, outliers) of AUC values at distances nearby or far from a small cell type from every random seed. High values represent a curve with greater divergence between near and far values indicating aggregation or repulsion, where lower values represent a more stable uniform distribution. **d**, **e**, Quantification of **c** comparing median values with or without the presence of large cells for the uniform (**d**) or grouped (**e**) distributions.

Using these layouts, we compared results from ClumPyCells with and without the size correction feature, alongside results from commonly used methods from spatstat^19^ and Squidpy.^18^ To measure the performance of each method, we reasoned that when observing aggregation of the two small cell types together, the uniform layout should have a flatter curve, while the grouped layout should have a curve with low values nearby and higher values farther away. To quantify the results of all random experiments, we compared the nearby and far distances by the area under the curve (AUC) of each method’s normalized output curve (Figures 2b and 2c).

Without large cells present, we expected the AUC difference within a cell type’s aggregation to be small in the uniform layout, indicating that a cell type was randomly distributed relative to itself (Figure 2c). In the grouped layout, we anticipated repulsion at near distances and a shift towards aggregation at further distances, resulting in a larger AUC difference. We observed this behavior in both ClumPyCells and spatstat. We expected our new implementation of the mark correlation function to be comparable to the equivalent method used by spatstat. Squidpy showed similar performance between the uniform and grouped layouts; upon further inspection of the output curves we observed erratic values with outliers driving the normalization resulting in reduced performance (Figure 2b).

To determine the effect of adding a few (15) large disruptions, we added large cells to the previous layouts(Figures 2d and 2e). As the underlying cell distributions were identical, we measured performance by comparing the median AUC difference of within-cell-type aggregation in each layout with and without large cells (Figures 2d and 2e). Across the uniform layouts, ClumPyCells was the most robust and interpretable with the addition of large cells. Although Squidpy appeared to stay consistent with large cell additions, close inspection revealed erratic and small values that collectively result in difficult to interpret results. Across the grouped layouts, ClumPyCells had the most stable median values. This relationship was also consistent across a mirrored analysis (Supplementary Figure S1). Together, these results demonstrate the reliability of ClumPyCells in the presence of large cells while maintaining correlation function capability.

### ClumPyCells identifies different scales of aggregation in treated melanoma

Given ClumPyCells’ ability to detect accurate cell type aggregation, we further assessed our method on biological samples. We applied ClumPyCells to a melanoma dataset consisting of 72 samples measured with IMC grouped into two cohorts.^27^ The first cohort contained 30 pretreatment samples from patients who received immune checkpoint inhibitor (ICI) treatments. Within these 30 samples, 14 patients demonstrated a response to ICI treatment, while the others were considered non-responders. The second cohort consists of 42 samples from ICI-untreated patients, including 5 benign nevi and 37 melanoma patients. Moldoveanu et al.^27^ quantified the tendency for each cell type to cluster. In these samples, melanoma cells, macrophages, monocytes, and “others” showed the highest level of aggregation, while regulatory T cells had the lowest assortativity coefficient. Using ClumPyCells, we found a similar relationship as melanoma cells had the highest aggregation across all samples, followed by macrophage, monocytes, and others, while regulatory T cells had the highest repulsion to itself (Figure 3a). Additionally, ClumPyCells found T and B cells agregate among themselves with a slight repulsion to other cell types, aligning with the previous analysis (Figure 3a).

**Figure 3:**
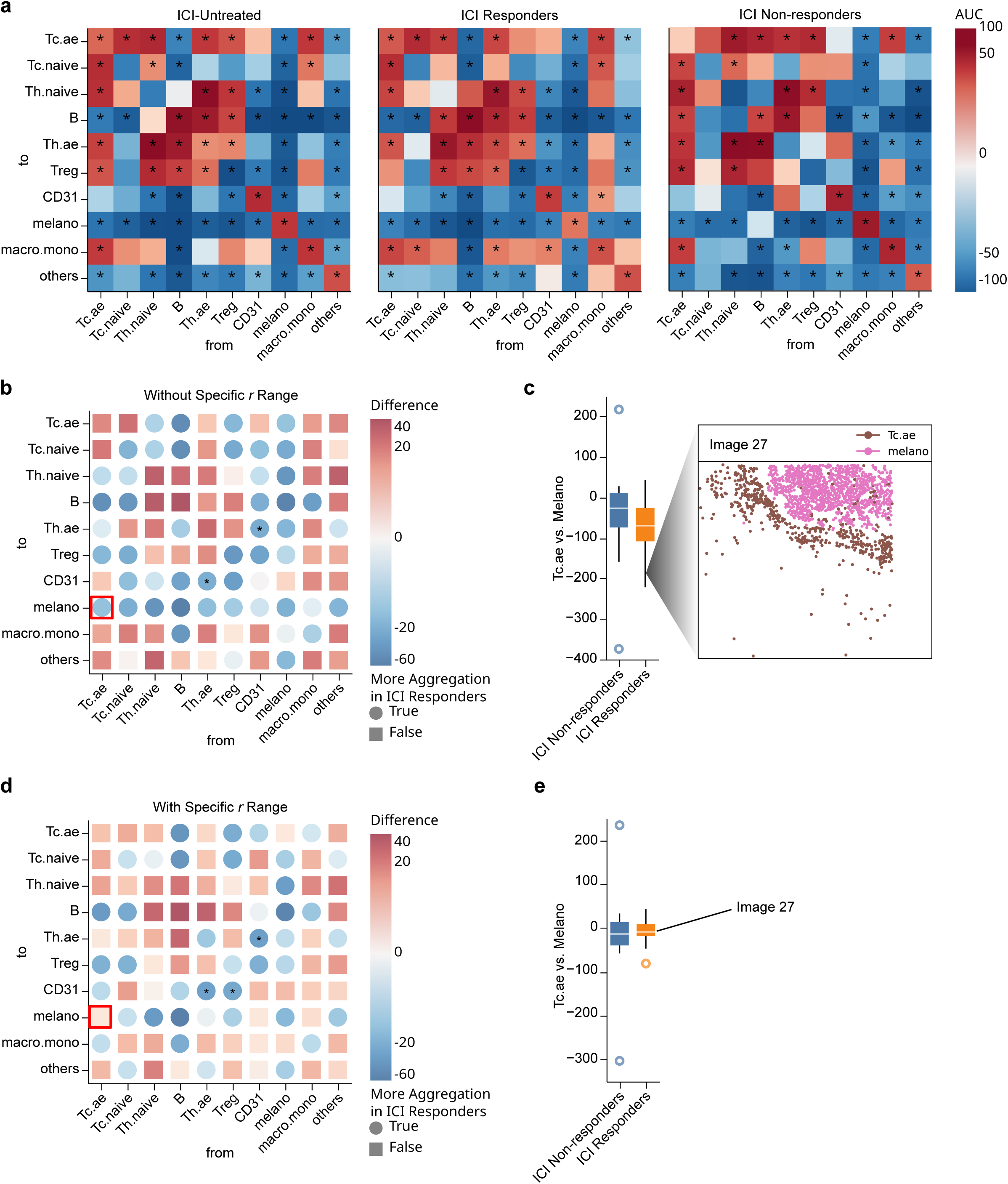
ClumPyCells identifies multi-resolution spatial relationships in melanoma samples from ICI-treated and ICI-untreated patients. **a**, Heatmaps of the mean ClumPyCells AUC values showing spatial relationships between different cell types in ICI-untreated patients (left), ICI responders (middle), and ICI non-responders (right; red: aggregation, blue: repulsion, *: two-sided permutation test, *p* 0.05). **b**, Heatmap showing the difference between ICI responders’ and ICI non-responders’ values from **a** (red square: higher aggregation in responders compared with non-responders, blue circle: higher repulsion in responders than non-responders, red box: aggregation value between melanoma and antigen-experience T cells, *: Fisher-Pitman permutation test, *p* 0.05). **c**, Left: Box-and-whisker plot showing the distribution of AUC values for spatial correlation between antigen-experienced T cells and melanoma cells in images from responders and non-responders (center line, median; box limits, upper (75^th^) and lower (25^th^) percentiles; whiskers, 1.5 × interquartile range). Right: Example image in the lower quartile for responders showing repulsion between antigen-experienced T cells and melanoma cells after ClumPyCells calculation but previously annotated as close proximity.^27^ **d**, **e**, As in **b**, **c**, but with the range of *r* for spatial correlation calculation restricted to values around median distance between focused cell types 120 *r* 150. Melanoma aggregation with antigen-experience T cells is now higher in responders (red box), and Image 27 is now ranked higher among responders. Nomenclature from Moldoveanu et al.,^27^ with antigen experienced (Tc.ae), naive (Tc.naive), and regulatory (Treg) T cells, naive (Th.naive) and antigen experienced (Th.ae) T helper cells, and macrophages and monocytes (macro.mono).

We further checked the correlation of T cells and cancer cells. Moldoveanu et al. reported that antigen-experienced T cells were located closer to cancer cells in pretreatment samples of responders than non-responders. However, our initial results showed slightly more repulsion between these two cell types (Figure 3b). Upon further investigation, we discovered that this discrepancy was due to the difference in the distance range being analyzed. Moldoveanu et al. used the median distance between two cell types, while ClumPyCells is based on correlation against random distribution. Thus, in cases where two cell types were close but clearly separated into clusters, their method considered the two cell types aggregated, whereas ClumPyCells identified them as self-aggregating but repulsive to the other cell type (Figure 3c). To resolve the discrepancy, we adjusted the distance range of the mark correlation function to be close to the median intercellular distance. After doing so, our results again aligned with the original finding, suggesting that ClumPyCells can find new biology based on the resolution required on the tissue (Figure 3d, Figure 3e). This analysis demonstrates the flexibility of ClumPyCells: the default range in ClumPyCells is appropriate for analyzing intermixed cells, while it can be applied to spatial relationships between clusters of the same cell type as well by setting the range close to the median distance.

### ClumPyCells classifies patients with AML from spatial aggregation

The bone marrow (BM) is a complex tissue in which hematopoietic stem and progenitor cells (HSPCs) differentiate into erythroid, myeloid, and lymphoid cell lineages. This process is tightly regulated by the hematopoietic microenvironment, which consists of stromal cells such as mesenchymal stromal cells (MSCs), adipocytes, megakaryocytes, and trabecular bone.^28–30^ Acute myeloid leukemia (AML) is an aggressive blood cancer that can disrupt the hematopoietic microenvironment, altering the distribution of cells and affecting their interactions. This disruption may lead to changes in cell aggregation patterns, which influences cell interactions and ultimately cell function. Accurately measuring the distribution of these cells, whether through cell-type or marker-based aggregation, is challenging. This difficulty arises from the variability in cell shapes and sizes, which creates unique spatial patterns in the BM, particularly those influenced by adipocytes. To map the spatial patterns of cells at single cell resolution while correcting for the influence of large adipocyte cells, we applied ClumPyCells to a data set of 36 subregions of interest (sub-ROIs) of BM from patients with AML (*n* = 12) and 15 sub-ROIs of normal BM (NBM; *n* = 5) imaged with IMC (Figure 4a, Supplementary Figures S2 and S3, and Supplementary Table S1). In NBM samples, CD34^+^ HSPCs were found to be primarily associated with erythroid cells (proerythroblasts; ECAD^+^; Figure 4a). However, this association may not be specific, as ECAD^+^ cells demonstrated interactions with all the assessed cell markers. HSPC showed repulsion among themselves (two-sided permutation test: *p* 0 01). This agreed with reports on mouse hematopoietic stem cells^31^ and with our own previous observations in human BM^32^ (Figure 4b), where HSPCs do not associate with each other. In AML, CD34^+^ cells exhibited only a trend toward self-repulsion. In NBM, CD3^+^ T cells aggregated strongly with CD20^+^ B cells, suggesting the presence of lymphoid aggregates^33^ (two-sided permutation test: *p* 0 01). Despite elevated blast counts, these associations persist in AML, suggesting remnants of an immune response (two-sided permutation test: *p* 0 01). Within NBM samples, ECAD^+^ cells significantly aggregated with all other cell the suppression of erythropoiesis disrupts ECAD aggregation to other immune cell types (Figure 4a). We found significant repulsion from MPO^+^ myeloid cells to adipocytes in NBM samples (two-sided permutation test: *p* 0 01), that may be explained by the increased density of MPO^+^ in these samples.^32^ In AML, MPO^+^ cells significantly aggregate with themselves, as well as with previously described blast markers such as CD68 (two-sided permutation test: *p* 0 01). Moreover, MPO^+^ cells also associate with Ki67^+^ proliferating cells, potentially due to the abundance of blasts that may also express MPO, ultimately affecting the distribution of all cell types.

**Figure 4:**
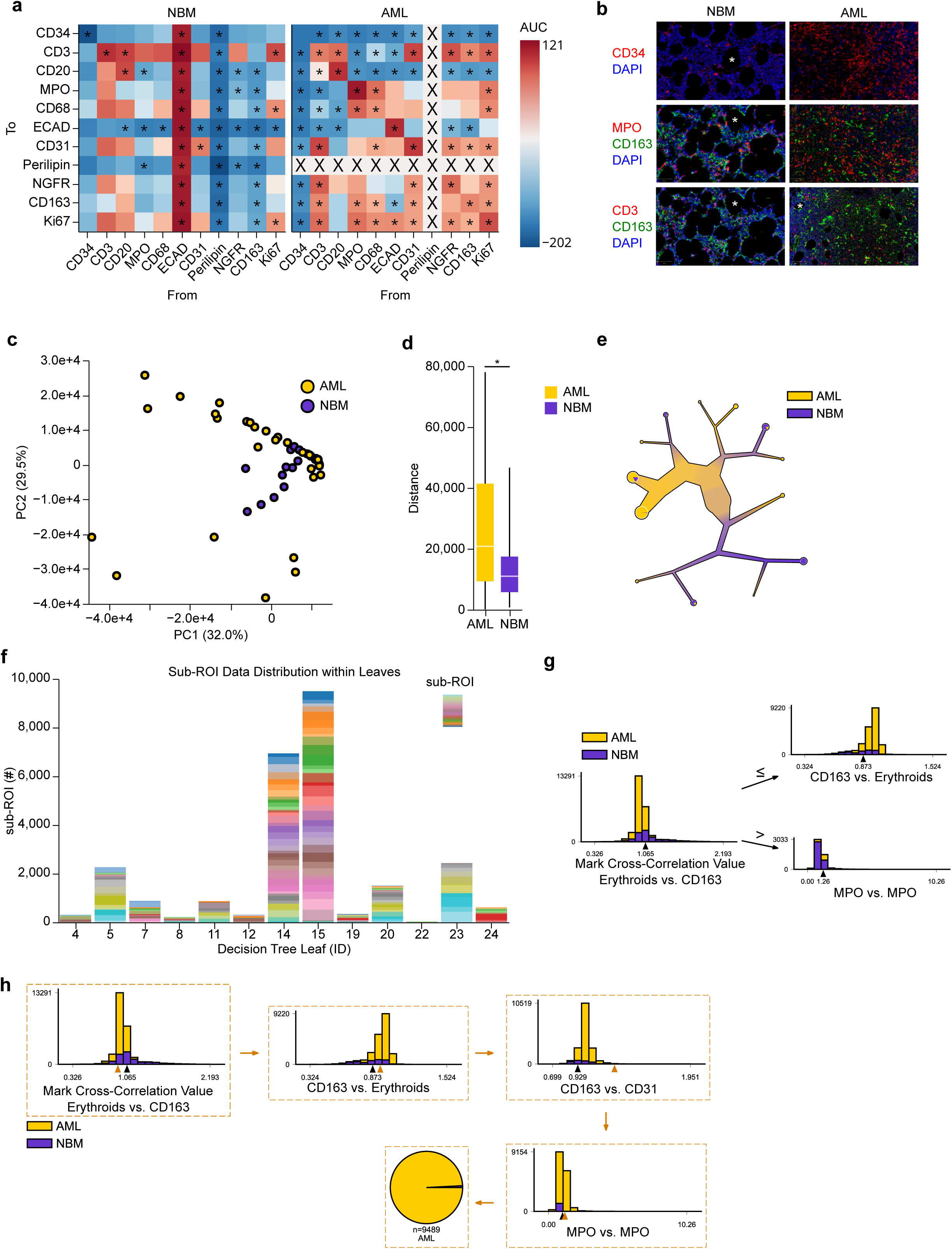
ClumPyCells classifies patients with AML using spatial correlations. **a**, Heatmaps of the mean ClumPyCells AUC values of the spatial relationship between different anti-body marker pairs in patients with AML (right) or normal bone marrow (NBM, left). Red: aggregation, blue: repulsion, “X”: too few cells, *: two-sided permutation test, *p* 0 05. **b**, Examples of marker expression in NBM and AML samples. Empty round spaces (asterisk) indicate adipocytes.**c**, Principal component analysis (PCA) of AML and NBM spatial correlations. **d**, Box-and-whisker plots (center line, median; box limits, upper (75^th^) and lower (25^th^) percentiles; whiskers, 1.5 interquartile range) of the pairwise distances in the PCA embedding from **c**. **e**, The full decision tree visualized with TooManyCells,^53^ with all observations starting in the root node (largest center circle) and ending in leaf nodes visualized with pie charts. **f**, Bar plot of the number of observations (*y*-axis) in each leaf node (*x*-axis), colored by sub-ROI. There is no leaf node dominated by a single sub-ROI. **g**, Bar plots of the tree nodes closest to the root, representing the most impactful pairs of comparisons. Each bar plot represents a node in the tree split by the number of aggregation observations (*y*-axis) from AML and NBM sub-ROIs. **h**, Path in the decision tree from the root node to the leaf node with the most AML-associated observations, with bar plots as in **g**.

Regarding stromal cells, such as NGFR^+^ MSCs, CD163^+^ macrophages, and adipocytes, we found significant repulsion to other stromal and hematopoietc cell types in NBM samples (two-sided permutation test: *p* 0 05), while conversely in AML samples there was increased aggregation (Figure 4a). These findings suggest that the stroma is evenly distributed in NBM, while in AML the stroma forms distinct cold areas without aggregation and hot areas characterized by stromal aggregation. There is reduced adipocyte abundance during AML development^34^ (Figure 4b), which may influence aggregation patterns with other cell types. NGFR^+^ MSCs and CD163^+^ macrophages acquire significant aggregation with CD3^+^ T cells and repulsion to CD20^+^ B cells (two-sided permutation test: *p* 0 01). MSCs and macrophages play an active role in lymphoid cell function and regulation (Figure 4a), and this role may still be active in AML.^35^

Leukemic diagnosis relies on the percentage of blasts in the BM. Although AML diagnosis starts when a patient’s blast percentage is at least 20%, there is a wide range of blast burdens. We reasoned that spatial pattern dynamics would differ between samples with different blast burdens. We stratified patients with 20-50% (low/intermediate blast count) and >60% blast count (high blast count) AML patients (Supplementary Figure S4). Cell proliferation and stromal cell markers were significantly more grouped together in high blast burden patients (Mann-Whitney *U* test: *p* 0 0208). This aligns with our previous observation that NGFR was more aggregated in AML, and confirms that aggregation of NGFR^+^ stromal cells could be a distinct feature of AML. In contrast, CD20^+^ B cells and CD163^+^ macrophages were more aggregated with CD31^+^ megakaryocytes in samples from patients with low blast percentages (Mann-Whitney *U* test: all *p* 0 05). These associations may indicate the remnants of normal hematopoiesis in the BM of low-blast-percentage AML patients, demonstrating that ClumPyCells can capture these distinct cell correlations within the BM.

To visualize more complex differences in cell type correlations for each patient, we applied principal component analysis (PCA) to all ClumPyCells outputs corrected for size (Figure 4c). Interestingly, sub-ROIs from patients with AML were more spread across the embedded space than sub-ROIs from patients with NBM, which tended to group together (Mann-Whitney *U* test: *p* = 3.19 × 10^−9^; Figure 4d and Supplementary Figure S5). This drastic change in variability suggests that BM in patients with AML has a greater diversity of spatial patterns than the normal BM niche, which may be driven by the percentage of blasts in the bone marrow.^36^

Motivated by this distinct difference between patients with and without AML, we developed a disease classifier using a decision tree based solely on the spatial pattern values found by our size-corrected ClumPyCells (Figure 4e and Supplementary Figure S6). After hyperparameter optimization, we found our decision tree model could achieve 86.5% accuracy in classifying disease state. Importantly, no single tree leaf node was influenced by a specific patient, which confirms a well-distributed tree (Figure 4f). We then examined the most distinct separator at the top-level decision, which differentiates the maximum number of points between AML and NBM. These nodes contain three pairs of correlations: erythroids and CD163^+^ macrophages, macrophages and erythroids, and MPO^+^ myeloids with themselves (Figure 4g). The top-most node of the tree may be explained by the influence of BM-derived macrophages on the formation of erythroid colonies.^37^ In AML, the disruption of erythroblastic islands and reprogramming of macrophages, including CD163^+^ subsets, can alter their proximity to erythroids, leading to measurable changes in spatial correlation patterns. Notably, CD163 is an M2 macrophage surface marker and is linked to poor prognosis.^38,39^ The path with the node collecting the largest number of observations from AML samples contained most of the top-level comparisons, acting as additional validation of the tree (Figure 4h).

To determine whether ClumPyCell marker correlations associate with patient survival outcomes, we used spatial correlation values as a covariate into a Cox proportional hazards regression (Supplementary Table S4). We found that CD34 vs. MPO was significantly associated with overall survival (Wald test: *p* 0 0328), highlighting the prognostic value of spatial marker relationships in differentiating AML from NBM. CD34 and MPO are two well known markers that impact patient outcome. CD34 expression is linked with lower complete response in de novo AML and is a poor prognostic factor. Conversely, MPO expression is associated with favorable karyotypes and an overall better prognosis. Together, these results suggest that size-corrected spatial correlation values not only sufficiently separate AML tissue from NBM, but may also predict patient outcome.

### Spatial transcriptomics analysis with ClumPyCells shows difference in tissue regions

ClumPyCells’ size-correction capability is flexible, enabling the exclusion of unwanted regions of tissue. To demonstrate this unique aspect, we applied ClumPyCells to colorec-tal cancer tissue measured with the recent Visium HD technology, providing single-cell resolution spatial transcriptomics. Characterizing the tumor microenvironment is crucial for understanding colon cancer progression and metastasis.^40^ However, the cell and gene distributions within different regions around the tumor are not fully understood. As such, we applied ClumPyCells to better map the cell niches within colon cancer tissue.

With expert annotation from a pathologist, we labeled five separate regions: normal colon, submucosa, adenoma, infiltrating tumor, and tumor boundary from a publicly available dataset of human colon cancer that contains normal and tumoral areas (Figure 5a). We hypothesized that the spatial distribution of markers among three cell populations (stroma, endothelial, and immune cells) would vary among regions. However, these populations may be artificially aggregated based on tumor or goblet cells, which force the other cells together in the tissue where they may still otherwise be randomly distributed. Therefore, in the tumor regions, we used ClumPyCells to remove pixels belonging to the previously-identified tumor clusters and goblet cells in the normal colon region. We found that the closer spatial association between the three populations was present in the tumor boundary, followed by the adenoma area and normal colon, where stromal and endothelial cells were spatially aggregated (Figure 5b). In contrast, the infiltrating tumor region showed a mild repulsion between stromal and endothelial cell types. The submucosa region, however, did not show any strong spatial association. As expected, immune cells tended to aggregate with each other at the tumor edge and infiltrating tumor regions.

**Figure 5:**
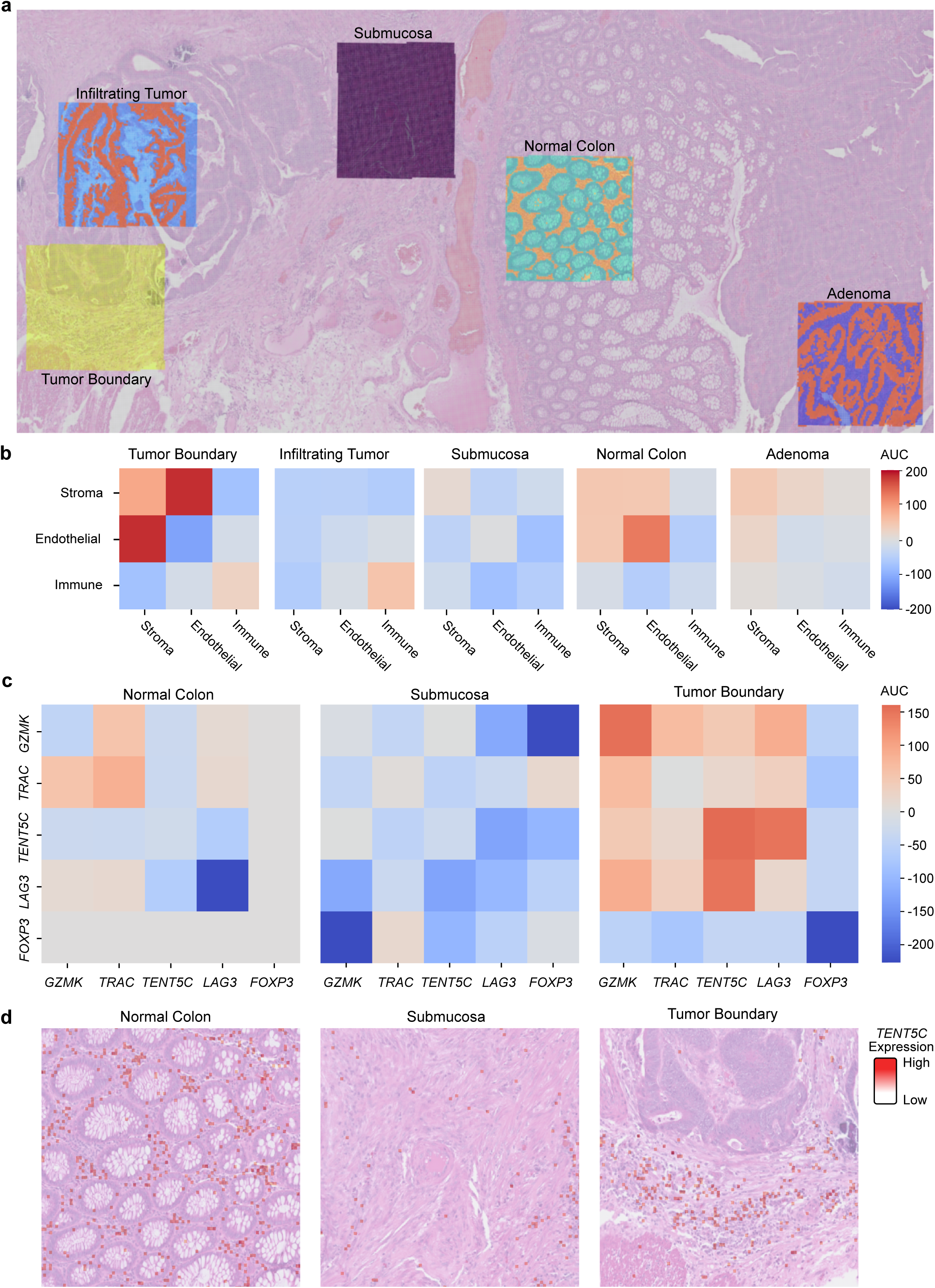
ClumPyCells analysis on Visium HD colorectal cancer data identifies T-cell aggregation patterns. **a**, Regions of interests (ROI) selected in an hematoxylin and eosin stained tissue with colon cancer. ROIs were labeled by a pathologist. **b**, ClumPyCells results measuring aggregation of different cell-type markers present in 8 µm squares in each ROI. In the tumor ROIs, cancer clusters were “removed” using the size correction feature of ClumPyCells, while in the normal colon region, goblet cell clusters were removed to determine true spatial distributions of other marker types (see Methods). Infiltrating tumor and tumor boundary ROIs displayed higher immune cell aggregation. **c**, ClumPyCells analysis specific to T cell biomarkers in each ROI as in **b**. **d** *TENT5C* expression overlayed in different ROIs. Although there is seemingly high aggregation in the normal colon, this is due to the presence of large structures (goblet cells), while the space between the goblet cells has a more uniform distribution.

As there were interesting spatial patterns of immune cells nearby tumor regions, we further investigated different immune-cell markers aggregating at these areas belonging to T cells, B cells, granulocytes, neutrophils, macrophages, and dendritic cells (Supplementary Figure S7). While we found that in the normal colon region there were associations between T cells and dendritic cells, there were large groups of different T-cell markers aggregated within the infiltrating tumor and tumor boundary (Figure 5c). The tumor boundary contained the highest aggregation among different cell marker pairs, such as B cells and T cells, and B cells and dendritic cells. Further investigation pointed to specific immune population markers (*GZMK*, *TENT5C*, and *LAG3*) aggregated in the tumor boundary, but with minimal aggregation elsewhere. More specifically, ClumPyCells identified *GZMK*, as a T-cell marker, more highly aggregated in the tumor boundary than normal colon.^41,42^ *GZMK* effector T cells are more enriched in glioblastoma, and similar tumor-infiltrating lymphocytes are present in various cancer types.^43^ Another T-cell biomarker, *TENT5C*, had higher aggregation in the tumor boundary region compared to all the other regions which may be due to its tumor suppressor function, as such genes are exected to be elevated in tumor regions^44–46^ (Figures 5c and 5d and Supplementary Figure S8). Together, these results demonstrate ClumPyCells’ capabilities to quantify immune aggregates within specific tissue regions. As gene signatures of tumor-infiltrating immune cells associate with prognosis,^47^ our observations suggest that ClumPyCells may contribute to prognostic predictions.

## Discussion

While current imaging technologies enable high-resolution analysis at the cellular level, it is crucial to account for biases introduced by cell morphology and tissue structure that may influence research findings. We observed significant differences in the interpretation of cell aggregation based on cell size. Considering the range of cell sizes within a sample, which can vary by an order of magnitude in diameter, we sought to overcome limitations of cell spatial correlation analysis by developing ClumPyCells, a new cell aggregation method which corrects for cell size. ClumPyCells not only provides size-corrected spatial relationship measurements between cells, but also between molecular expressions tested across several technologies, including IMC for protein abundance and Visium HD for subcellular gene expression. ClumPyCells also provides a new permutation test for spatial analysis, ensuring confidence in the reported results. ClumpPyCells is a flexible library that fully integrates with SquidPy.

To observe the effect of large cell size disruption within a tissue as well as benchmark and compare ClumPyCells with popular alternative methods without size correction capabilities, we simulated cell distributions using different cell sizes. We found that ClumpyCells outperformed other methods by presenting smoother correlation functions with lower impact from the introduction of large cells. This result suggests that ClumPyCells takes the necessary aspects from each of the other methods, including smoothing from spatstat and the potentially lower variability from SquidPy.

When applied to samples from a melanoma data set, we observed the advantage of another desired aspect of ClumPyCells: multi-resolution aggregation. While at close distances, T cells and cancer cells were more aggregated with themselves, at larger distances they were close to each other, suggesting that these cell types are generally close together within tissue, but in a self-aggregating pattern. This finding both reinforces and bolsters the original interpretation of the data set due to a multi-resolution interpretation. We also demonstrated the use of spatial analysis in disease prediction and overall survival. Using ClumPyCells to measure molecular aggregation within samples from a cohort of AML and control patients, we identified significant differences in the spatial proximity of proteins between conditions. We constructed a decision tree to characterize the importance of each aggregation and obtain a non-linear classification of differences between AML patients and healthy individuals. ClumPyCells reports variables that can directly be used in a proportional hazards model to determine effects on overall survival.

A current limitation for training the decision tree on the AML cohort is the use of mark correlation values across all distances from origin cells to help offset the limited data size. Based on our results, we note that observations at specific distance ranges may contribute more to differentiating between AML patients and healthy individuals. We propose with larger cohorts to use clustering techniques to identify a range of distances that, across all images, will have the most significant impact on spatial correlation results. In addition, we expect that future studies with a large-scale cohort paired with fully annotated scanned samples for cell types will make full use of survival analyses, pinpointing cell types and tissue structures associated with survival.

Importantly, we also demonstrated that, in addition to cellular-level analysis in protein-based assays, ClumPyCells is flexible enough to use with gene expression. With recent spatial transcriptomic technologies such as Visium HD, our method was able to identify bio-logically relevant expression correlations in the colorectal cancer tumor microenvironment. This analysis is uniquely capable with ClumPyCells, purposefully “size correcting” tumor and goblet cells to focus solely on the distribution of molecular markers associated with other cell types such as immune cells. This flexible method, which works across technologies, data modalities, and scales, exemplifies the need for additional corrections for cell morphologies measured with these assays. As such, we anticipate that ClumPyCells will push bioimaging analysis forward and contribute to unraveling the tissue microenvironment.

## Methods

### ClumPyCells Aggregation Measurement

ClumPyCells’ foundations are built on point processing methods to analyze spatial relationships between different cell or marker types. Initially, we segment input images with CellProfiler.^24^ We represent these distributed points as a *c × m* matrix (encoded as a data frame with *x* and *y* coordinates) with *c* cells or spatial transcriptomic bin barcodes and *m* “marks”. is an additional matrix containing other relevant information, such as continuous variables (e.g. cell radius, cell area, intensity of a fluorescence markers) and discrete variables (e.g. cell type). ClumPyCells measures the aggregation between different “marks” using the mark cross-correlation function,

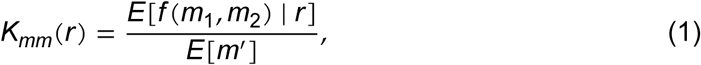

where *K_mm_* (*r*) measures the expected value of a function *f* of two marks *m*_1_ and *m*_2_ at a distance *r*, normalized by the expected value of the marginal distribution of the specified mark *m*^′^. This function allows ClumPyCells to quantify the degree of co-localization or segregation between different cell types or features within a specified spatial range.

The mark correlation function *K_mm_* (*r*) is a function of *r*, which ranges from 0 to a user-specified maximum distance *r*_max_. The test function *f* is flexible, but here defined as

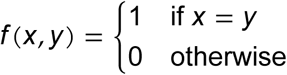

for discrete values or

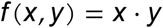

for continuous values.

The numerator of Equation (1) calculates the average amount of the mark at distance *r*, while the denominator calculates the average amount of the mark in the entire image. Therefore, *K_mm_* (*r*) > 1 indicates that more of the mark is found at distance *r* than would be expected by random distribution, suggesting aggregation. Conversely, *K_mm_* (*r*) < 1 indicates fewer marks at this distance than expected, suggesting repulsion. A value of *K_mm_* (*r*) = 1 indicates that the mark is randomly distributed at this distance.

### Size Correction Algorithm

In order to mitigate the effects of cells such as adipocytes occupying a proportionally larger space and forcing cells such as B cells and T cells towards each other, we designed a size correction algorithm for ClumPyCells. The numerator of the mark correlation function is the expected value of the tested amount of mark at a given distance *r*. The size correction algorithm only accounts for the intercellular distance, ignoring intracellular areas while calculating the distance. We define this corrected distance *r* between two cells *p*_1_ and *p*_2_ as

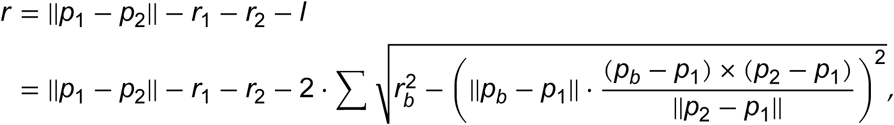

where *p_b_* is a large cell in-between *p*_1_ and *p*_2_. *l* is the length passing through the large cell. *r*_1_, *r*_2_, and *r_b_* are the radii for *p*_1_, *p*_2_, and *p_b_* respectively.

### Min-mid-max Normalization

Since the output of the mark correlation function consists of curves that vary across different distances, we must summarize each curve into a single value to represent the overall trend of aggregation or repulsion for simpler quantification. The most straightforward approach is to calculate the AUC relative to *K_mm_* = 1. This method involves converting correlation values where *K_mm_* > 1 as positive and areas where *K_mm_* < 1 as negative so that repulsion values negate aggregation values. However, since the range for aggregation is (1, ∞) while the range for repulsion is [0, 1), this method causes the aggregation part to dominate.

As the mark correlation function is a ratio, another approach is to transform the correlation curve with the logarithm, converting the range of aggregation to (0, ∞) and repulsion to (-∞, 0). However, we found that because the same type of cells are at certain distances are usually few, there is limited aggregation but much repulsion which results in many near-zero values since, at certain distances, no points with the same type can be found. This results in a dominating repulsion value.

Instead, we developed a new normalization for spatial correlation values called “min-mid-max” normalization. For each spatial correlation value for some *r*, we normalize between a maximum and middle value (here *K_mm_* = 1) to be 0 and 1, and separately normalize the middle and the minimum values (here *K_mm_* = 0) as 0 and −1. Then, for quantification, we sum all points for a final AUC to represent the entire curve, or specific *r* values to represent a region of the curve. We selected maximum and minimum values based on all values across all comparisons in all sub-ROIs. To prevent outlier values from skewing the results, we treated values that are three times the interquartile range away from the median value as outliers and selected the maximum and minimum values after excluding them.

We provide users with both the logarithm and the min-mid-max normalization methods in ClumPyCells. When there are many cells present but only a few types of cells, the logarithm method may be sufficient. Otherwise, the min-mid-max normalization represents the overall trend more accurately.

### Benchmarking Experiments

We designed four types of simulated data sets, each with 100 randomly-generated synthetic tissues. Two of these layouts contained only two cell types of equal sizes, while the remaining two layouts contained a third cell type as well that was around 40 times larger than the other two cell types. The first type of layout, called “uniform”, drew cell types from a uniform distribution. The second layout, called “grouped”, was an aggregated distribution of the two smaller cell types with limited ranges at separate corners of the tissue. In contrast, the third and fourth layouts repeated these first two layouts but added large cells to represent adipocytes as in tissue from bone marrow.

For each layout, we compared the performance of four methods: ClumPyCells with size correction, ClumPyCells without size correction (a new implementation of a standard mark correlation function as in spatstat), spatstat, and the co-occurrence probability method from SquidPy. The outputs of the four methods included the radius of measurement and the correlation score for each radius. To compare the methods more effectively, we applied min-mid-max normalization to the output plots of all methods.

To quantify the performance of each method, we measured the AUC of two radius ranges: near and far. The first AUC ranged from 0 to 10 to quantify distribution for the area representing nearby cells, and the second AUC ranged from 150 to 160 (given the maximum tissue size was 330) to quantify the area far from each cell. We expected the absolute difference between the near and far AUC values to be large for the grouped layout case and small for the uniformly distributed layout.

### Survival Analysis

Using the mark correlation function results combined with clinical data as features, we analyzed the factors affecting patients’ survival with a Cox proportional hazards model.^48^ Due to the limited cohort size, we performed feature selection before training the model. In ClumPyCells, we proposed two ways of feature selection methods: selecting the features with the highest individual hazard coefficient or with the highest individual significance. After fitting the model, we identified the significant features.

### Melanoma Data Preprocessing

#### Statistical Test Definition for ClumPyCells

The summarized spatial relationship reported by ClumPyCells averages the AUC values from each sub-ROI, AUC (*K_mm_*). To determine if this value was significantly aggregated or repulsed, we developed a complementary statistical test for use with the (optionally size-corrected) mark correlation function. We designed a permutation test by constructing a null model of randomly shuffled marker labels within each sub-ROI, maintaining the spatial coordinates of each cell location. Before shuffling, we stored the large cell annotations so that the size correction algorithm would work as intended. Due to the computational complexity of calculating massive amounts of mark correlation functions, we shuffled the data 100 times, passing each new data set into ClumPyCells to get AUC (*K_mm_*). We compared the new permutation results with the results obtained from the original observed dataset, AUC (*K_mm_*)_obs_. This observed result was considered consistent with the null hypothesis if:

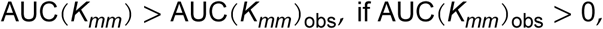

or

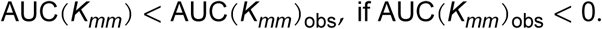

We then divided the number of permutations that supported the observation falling within the null hypothesis by the total number of permutations conducted, repeating this process for each correlation result.

### NBM and AML dataset

All biological samples were collected with informed consent in accordance with the Declaration of Helsinki and approved by either the University Health Network Research Ethics Board (02-0763) or the Instituto Mexicano del Seguro Social (R-2012-785-092). We recorded age, biological sex, and clinical information where available. We did not collect any data regarding ethnicity. We performed IMC staining and ablation as described previously.^49^ In-formation regarding antibodies resides in Supplementary Table S3. We generated overlaid images using Wolfram Mathematica v10.3 (Fluidigm), and we applied contrast enhancement using Adobe Photoshop solely for visualization purposes. We conducted image analysis and quantification following established protocols,^50,51^ with all data processed using standardized pipelines to ensure reproducibility.^52^ Our NBM and AML data set consists of IMC images of representative selected ROIs from myeloid biopsies of 12 AML patients and 5 patients with NBM. From each ROI, we further randomly selected three sub-ROIs of the same dimensions (1,056 px × 642 px) to avoid regions occupied by blood vessels and bones, resulting in 36 AML sub-ROIs and 15 NBM sub-ROIs. We segmented sub-ROI images with CellProfiler to obtain *x* and *y* coordinates, the mean intensity of each antibody marker, and the size of each cell.

### PCA Analysis

For each patient in the bone marrow cohort, ClumPyCells reported 121 correlation results of 11 cell markers. We used PCA with all 121 features and found that the first three principal components represented 75% of the variance in the data set. We determined overall distances between either sub-ROIs or patients by calculating pairwise distances between PCA observations.

### AML classifier

To further explore the nonlinear spatial relationship that distinguishes patient with AML from those without AML, we trained a decision tree classifier with ClumPyCells spatial correlation values as features and the image labeled by AML or NBM origin. Generated from 51 biopsies in total and 121 pairs of cell comparisons, the model was prone to overfitting. To address this, we performed data augmentation by treating the spatial correlation values between pairs of cell types at each radius as individual data points. This approach expanded the dataset to 26,163 data points, each with 121 spatial features.

To further mitigate overfitting, we carefully selected hyperparameters, including the maximum depth and the maximum number of features, through Bayesian optimization employed with 100 random initial points and 30 maximization steps. We evaluated the model based on test accuracy using three-fold cross-validation. The decision tree was visualized using TooManyCells^53^ for a holistic shape and dtreeviz for detailed statistics.

### Visium HD Annotation

We selected a publicly-available dataset of Visium HD, single-cell resolution spatial transcriptomics from a tissue with colorectal cancer to demonstrate the application of ClumPyCells to gene expression data. A certified pathologist divided the tissue into five spatially and histologically distinctive areas: (1) Normal colon mucosa: composed of the epithelium, lamina propria, and muscularis mucosa. The Epithelium is composed of a single layer of the columnar and goblet cells. (2) Submucosa: consists of loose connective tissue, smooth muscle bundles, vessels, and nerve plexuses. (3) Adenoma: the polypoid area, composed of dysplastic colorectal epithelium protruding into the lumen of the colon. This adenoma is a premalignant neoplasm confined to the superficial layers with no evidence of invasion. (4) Tumor boundary: the presence of single or small clusters of tumor cells (fewer than five cancer cells) at the invasive front of the tumor. (5) Tumor: an infiltrating tumor, moderately differentiated, with a morphology of gland forming carcinoma with desmoplasia. Glands are filled with necrotic debris (dirty necrosis) focally. We selected markers for immune cells present in colorectal cancer from the 10X Genomics Xenium Panel `hColon_V1`.

## Data Availability

The melanoma cancer dataset is available from https://doi.org/10.5281/zenodo.590 `3179`. Raw and annotated IMC bone marrow data is available at https://zenodo.org/records/14711407. Colorectal cancer Visium HD data is available from the 10X Genomics website (https://www.10xgenomics.com/datasets/visium-hd-cytassist-gene-expression-libraries-of-human-crc).

## Code Availability

ClumPyCells is available at https://github.com/schwartzlab-methods/ClumPyCells. Scripts for generating the figures and plots of the manuscript can be found at https://github.com/schwartzlab-methods/ClumPyCells_paper_figure.

## Acknowledgments

This work was supported by the Canadian Cancer Society (grant 707484; G. W. S.), the Natural Sciences and Engineering Research Council of Canada (grants RGPIN-2023-04713 and DGECR-2023-00395; G. W. S.), the Social Sciences and Humanities Research Council of Canada (grant NFRFE-2022-00681; G. W. S.), the Canada Research Chairs Program (grant CRC-2021-00240; G. W. S.), the Canadian Institutes of Health Research (grant PJT-203941; G. W. S.), the Princess Margaret Cancer Foundation (G. W. S.), and from CONACYT (grants CB-2012-179417, FC-205-2-1341_CCTyF; E. F.). This work was carried out with the aid of a grant from the International Development Research Centre, Ottawa, Canada (grant 109153-001; E. F.); the views expressed herein do not necessarily represent those of IDRC or its Board of Governors. The authors thank Dr. John Dick (University Health Network, Toronto, Canada), Olga Ornatsky (Fluidigm, Toronto, Canada), Dr. Ricardo Esquivel-Gómez (IMSS, Mexico City México), Dr. Antonio Hernández-Ramírez (IMSS, Mexico City México), Dr. Carlos R Hernández Pérez (IMSS, Mexico City México), and Dr. Eduardo Terreros (IMSS, Mexico City México). A. G. A. and E. F. received a scholarship from Programa de Cooperación Internacional del Instituto Mexicano del Seguro Social.

## Authors Contributions

G. W. S. and E. F. conceived and supervised the project. Z. Z. and J. C. developed the ClumPyCells method and software. Z. Z. developed and ran benchmarks. Z. Z. and J. C. ran and analyzed data. E. F. generated experimental results. Z. Z., J. C., A. G. A., E. F., and G. W. S. wrote and edited the manuscript. All authors reviewed the manuscript.

## Competing Interests

The authors declare no competing interests.

## Notes

https://github.com/schwartzlab-methods/ClumPyCells

### Competing Interest Statement

The authors have declared no competing interest.

