## Supplementary Information for "ClumPyCells resolves spatial aggregation in complex tissues overcoming size biases"

### Supplementary Figures

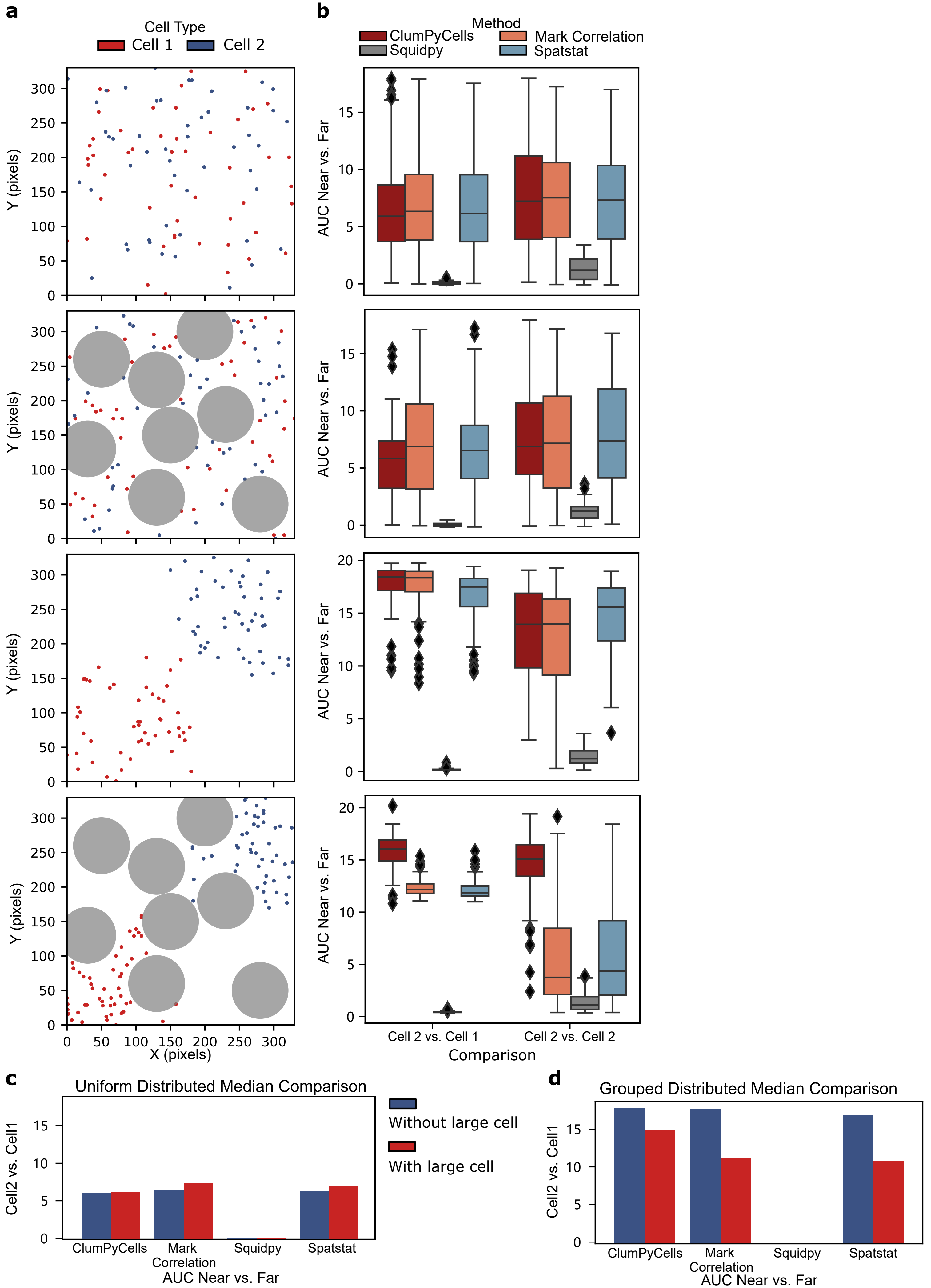
a
b
Cell Type
Cell 1
Cell 2
Method
ClumPyCells
Mark Correlation
Squidpy
Spatstat
Y (pixels)
X (pixels)
AUC Near vs. Far
Cell 2 vs. Cell 1
Cell 2 vs. Cell 2
Comparison
c
Uniform Distributed Median Comparison
Without large cell
With large cell
Cell2 vs. Cell1
ClumPyCells
Mark Correlation
Squidpy
Spatstat
AUC Near vs. Far
d
Grouped Distributed Median Comparison
Cell2 vs. Cell1
ClumPyCells
Mark Correlation
Squidpy
Spatstat
AUC Near vs. Far

Supplementary Figure S1: ClumPyCells is robust against large-cell presence in synthetic tissue benchmarks for remaining pairwise comparisons from Figure 2. **a**, **b**, As in Figures 2a and 2c, but comparing Cell 2 vs. Cell 1 and Cell 2 vs. Cell 2 ($n = 100$ for each layout, with box-and-whisker plots having center line, median; box limits, upper ($75^{th}$) and lower ($25^{th}$) percentiles; whiskers, $1.5 \times$ interquartile range; points, outliers). **c**, **d** As in Figures 2d and 2e, but quantification of **b**.

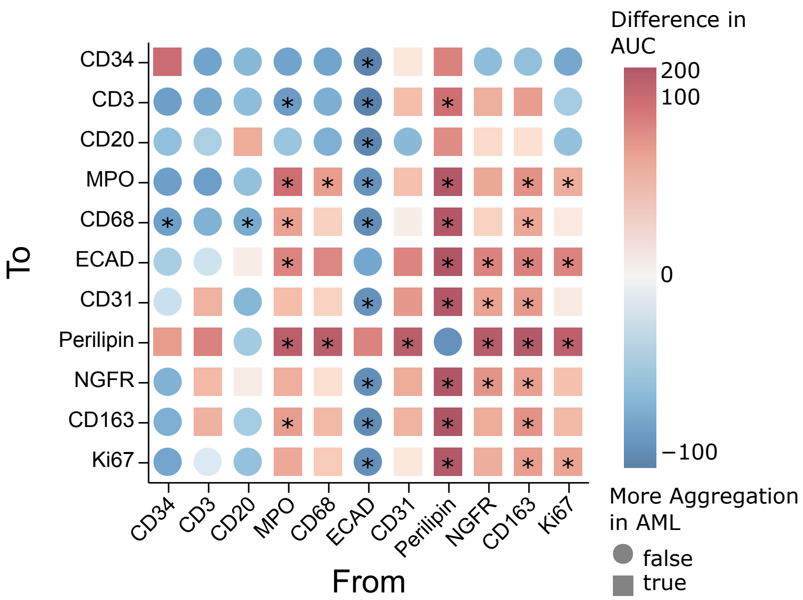

Difference in AUC
200
100
0
−100
More Aggregation in AML
false
true
To
CD34
CD3
CD20
MPO
CD68
ECAD
CD31
Perilipin
NGFR
CD163
Ki67
From

Supplementary Figure S2: Heatmap showing the difference between ClumPyCells AUC values between heatmaps generated from AML and NBM samples shown in Figure 4. Blue circle: more aggregation in NBM, red square: more aggregation in AML, *: Mann-Whitney $U$ test, $p < 0.05$.

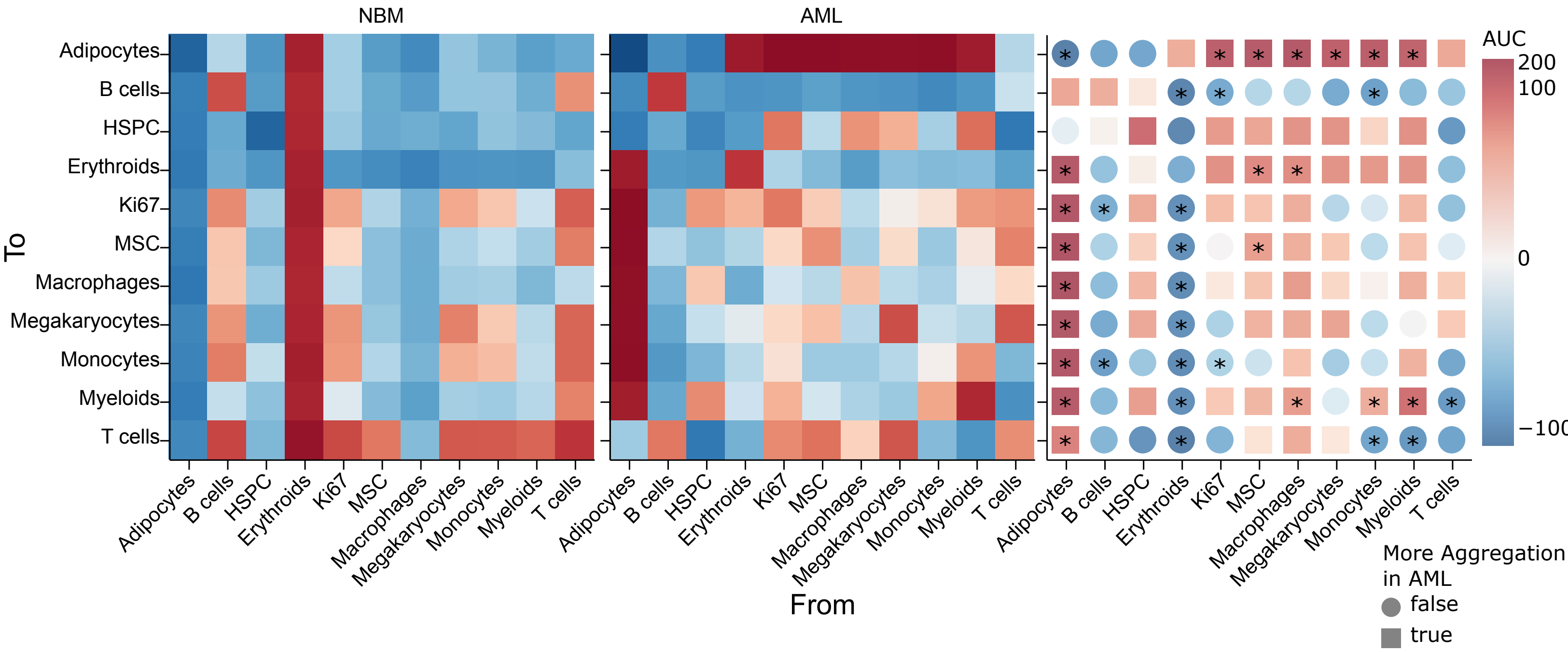

NBM
AML
To
From
Adipocytes
B cells
HSPC
Erythroids
Ki67
MSC
Macrophages
Megakaryocytes
Monocytes
Myeloids
T cells
AUC
200
100
0
−100
More Aggregation in AML
false
true

Supplementary Figure S3: Heatmaps of the mean ClumPyCells AUC values of the spatial relationship between different cell types in patients with AML (left) or normal bone marrow (NBM, middle), and the difference between both other heatmaps (right). Red: aggregation, blue: repulsion, circle: more aggregation in NBM, square: more aggregation in AML (*: two-sided permutation test, $p < 0.05$).

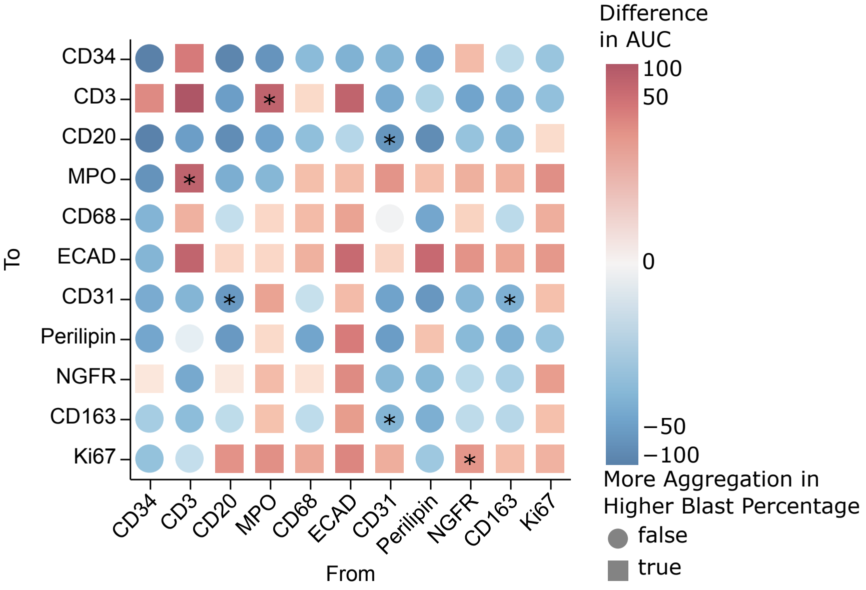

Difference
in AUC
100
50
0
−50
−100
To
CD34
CD3
CD20
MPO
CD68
ECAD
CD31
Perilipin
NGFR
CD163
Ki67
From
More Aggregation in
Higher Blast Percentage
false
true

Supplementary Figure S4: Heatmap showing the difference between AML patients with higher (greater than 60%) and lower blast percentage.

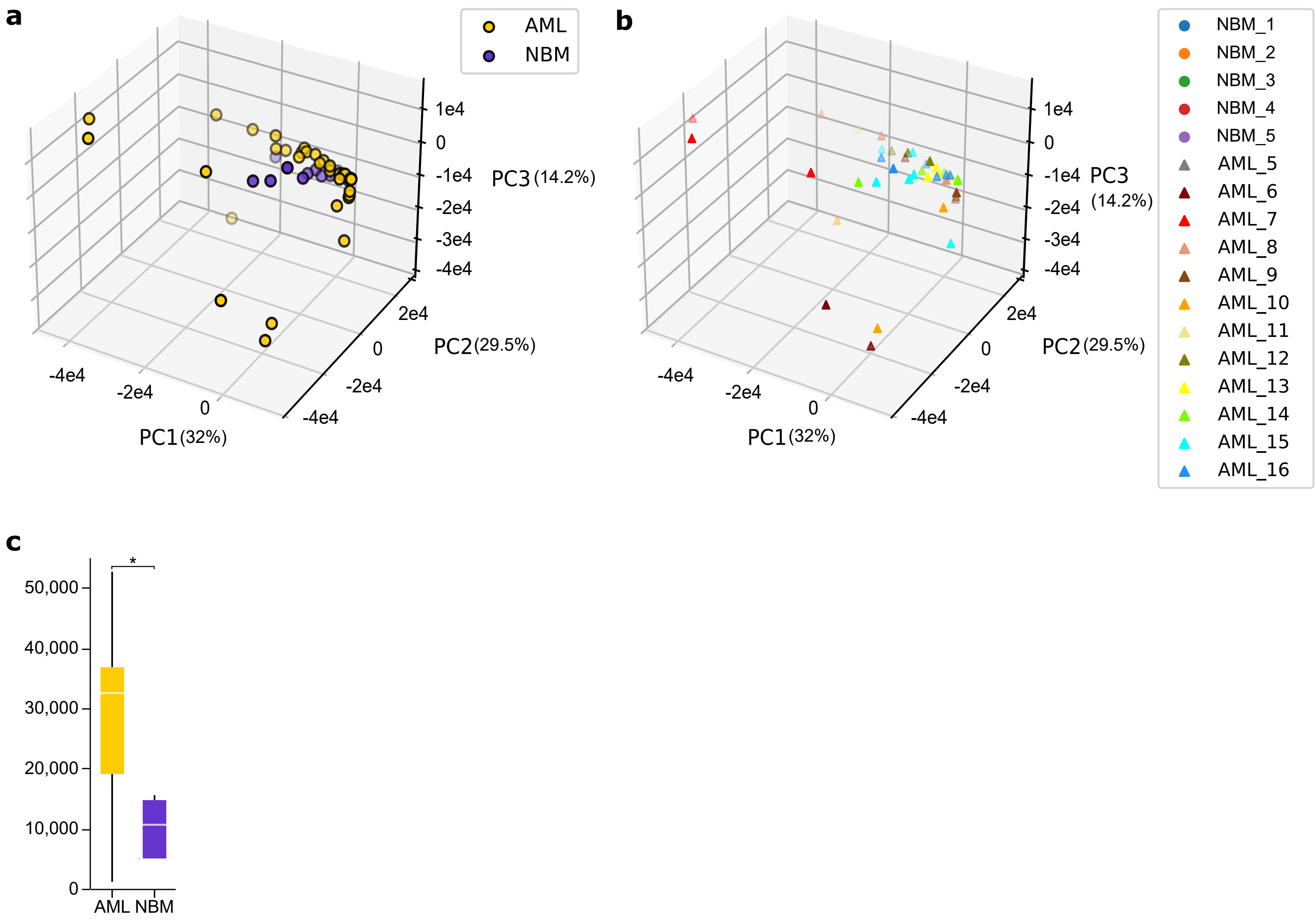
a
AML
NBM
1e4
0
-1e4
-2e4
-3e4
-4e4
PC3 (14.2%)
2e4
0
-2e4
-4e4
PC2 (29.5%)
-4e4
-2e4
0
PC1 (32%)
b
NBM_1
NBM_2
NBM_3
NBM_4
NBM_5
AML_5
AML_6
AML_7
AML_8
AML_9
AML_10
AML_11
AML_12
AML_13
AML_14
AML_15
AML_16
1e4
0
-1e4
-2e4
-3e4
-4e4
PC3 (14.2%)
2e4
0
-2e4
-4e4
PC2 (29.5%)
-4e4
-2e4
0
PC1 (32%)
c
*
50,000
40,000
30,000
20,000
10,000
0
AML
NBM

Supplementary Figure S5: No single patient is driving the variability between AML and NBM. **a**, **b**, 3D PCA of cell-type correlation values for each sub-ROI colored by condition (**a**) or patient with sub-ROI (**b**). **c**, Box-and-whisker plots (center line, median; box limits, upper (75th) and lower (25th) percentiles; whiskers, 1.5 $\times$ interquartile range) of the average pairwise distances within each patient in the PCA embedding from **b**.

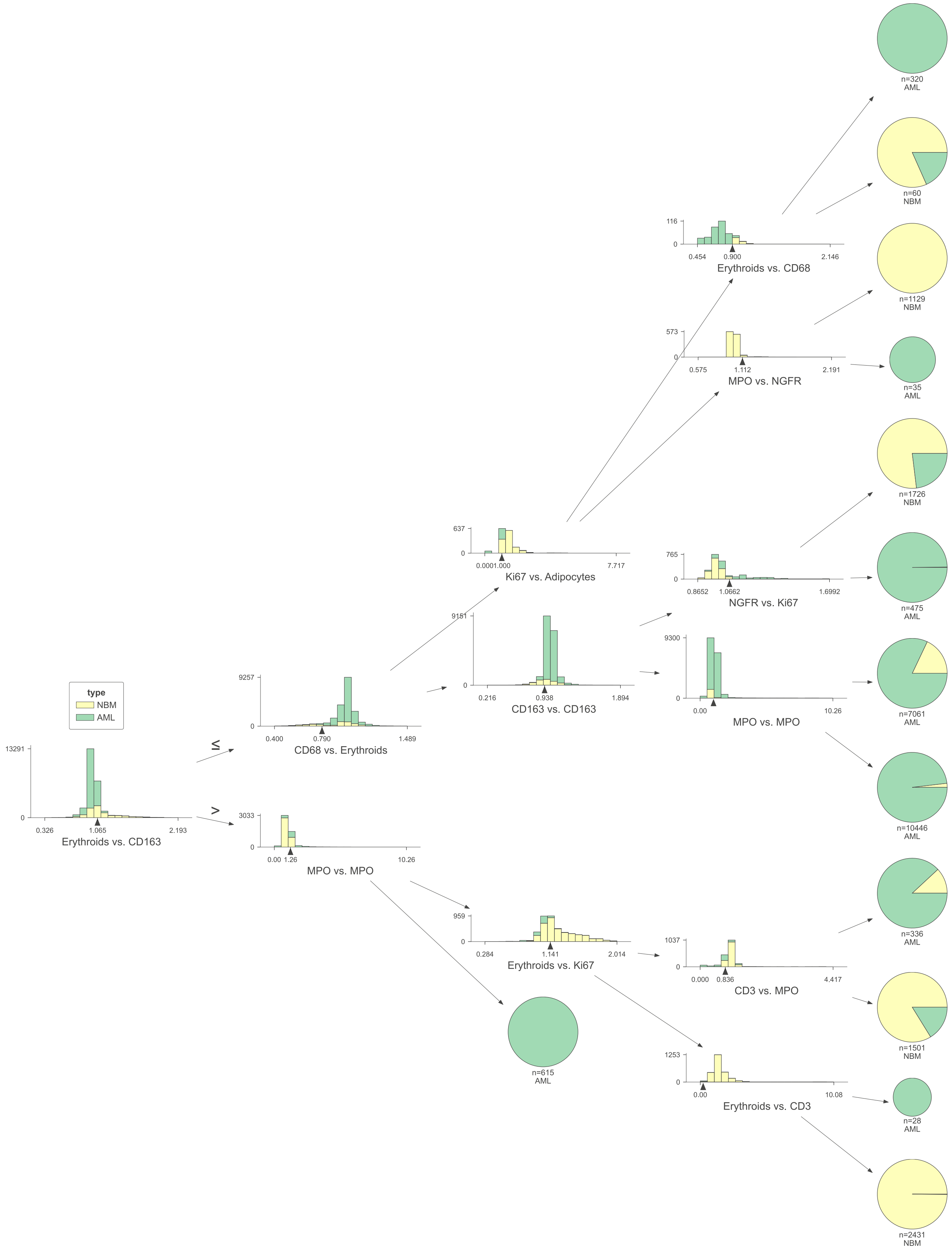

type
NBM
AML
13291
0
0.326
1.065
2.193
Erythroids vs. CD163
≤
>
9257
0
0.400
0.790
1.489
CD68 vs. Erythroids
3033
0
0.00
1.26
10.26
MPO vs. MPO
637
0
0.0001.000
7.717
Ki67 vs. Adipocytes
9151
0
0.216
0.938
1.894
CD163 vs. CD163
959
0
0.284
1.141
2.014
Erythroids vs. Ki67
n=615
AML
116
0
0.454
0.900
2.146
Erythroids vs. CD68
573
0
0.575
1.112
2.191
MPO vs. NGFR
765
0
0.8652
1.0662
1.6992
NGFR vs. Ki67
9300
0
0.00
10.26
MPO vs. MPO
1037
0
0.000
0.836
4.417
CD3 vs. MPO
1253
0
0.00
10.08
Erythroids vs. CD3
n=320
AML
n=60
NBM
n=1129
NBM
n=35
AML
n=1726
NBM
n=475
AML
n=7061
AML
n=10446
AML
n=336
AML
n=1501
NBM
n=28
AML
n=2431
NBM

Supplementary Figure S6: The complete disease classifier decision tree. Each leaf node represents a group of cell spatial correlation measurement.

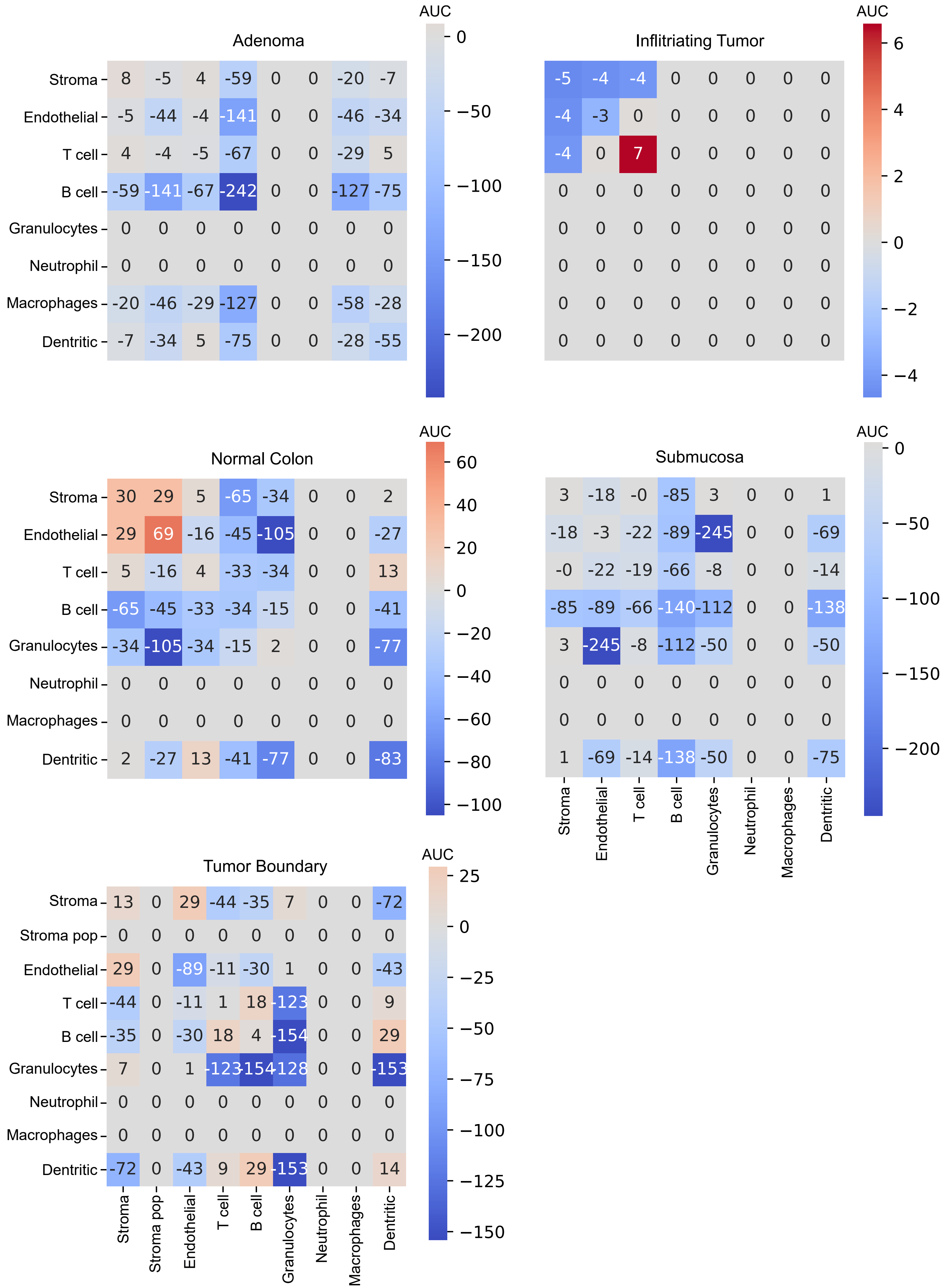

Adenoma
AUC
0
−50
−100
−150
−200
Stroma 8 -5 4 -59 0 0 -20 -7
Endothelial -5 -44 -4 -141 0 0 -46 -34
T cell 4 -4 -5 -67 0 0 -29 5
B cell -59 -141 -67 -242 0 0 -127 -75
Granulocytes 0 0 0 0 0 0 0 0
Neutrophil 0 0 0 0 0 0 0 0
Macrophages -20 -46 -29 -127 0 0 -58 -28
Dentritic -7 -34 5 -75 0 0 -28 -55
Inflitriating Tumor
AUC
6
4
2
0
−2
−4
-5 -4 -4 0 0 0 0 0
-4 -3 0 0 0 0 0 0
-4 0 7 0 0 0 0 0
0 0 0 0 0 0 0 0
0 0 0 0 0 0 0 0
0 0 0 0 0 0 0 0
0 0 0 0 0 0 0 0
0 0 0 0 0 0 0 0
Normal Colon
AUC
60
40
20
0
−20
−40
−60
−80
−100
Stroma 30 29 5 -65 -34 0 0 2
Endothelial 29 69 -16 -45 -105 0 0 -27
T cell 5 -16 4 -33 -34 0 0 13
B cell -65 -45 -33 -34 -15 0 0 -41
Granulocytes -34 -105 -34 -15 2 0 0 -77
Neutrophil 0 0 0 0 0 0 0 0
Macrophages 0 0 0 0 0 0 0 0
Dentritic 2 -27 13 -41 -77 0 0 -83
Submucosa
AUC
0
−50
−100
−150
−200
3 -18 -0 -85 3 0 0 1
-18 -3 -22 -89 -245 0 0 -69
-0 -22 -19 -66 -8 0 0 -14
-85 -89 -66 -140 -112 0 0 -138
3 -245 -8 -112 -50 0 0 -50
0 0 0 0 0 0 0 0
0 0 0 0 0 0 0 0
1 -69 -14 -138 -50 0 0 -75
Stroma Endothelial T cell B cell Granulocytes Neutrophil Macrophages Dentritic
Tumor Boundary
AUC
25
0
−25
−50
−75
−100
−125
−150
Stroma 13 0 29 -44 -35 7 0 0 -72
Stroma pop 0 0 0 0 0 0 0 0 0
Endothelial 29 0 -89 -11 -30 1 0 0 -43
T cell -44 0 -11 1 18 -123 0 0 9
B cell -35 0 -30 18 4 -154 0 0 29
Granulocytes 7 0 1 -123 -154 -128 0 0 -153
Neutrophil 0 0 0 0 0 0 0 0 0
Macrophages 0 0 0 0 0 0 0 0 0
Dentritic -72 0 -43 9 29 -153 0 0 14
Stroma Stroma pop Endothelial T cell B cell Granulocytes Neutrophil Macrophages Dentritic

Supplementary Figure S7: Cell-type aggregation measurements in a sample of colorectal cancer measured with Visium HD from Figure 5a in different regions of interests. Red: aggregation, blue: repulsion, gray 0 value: no neighboring cells of the specified type.

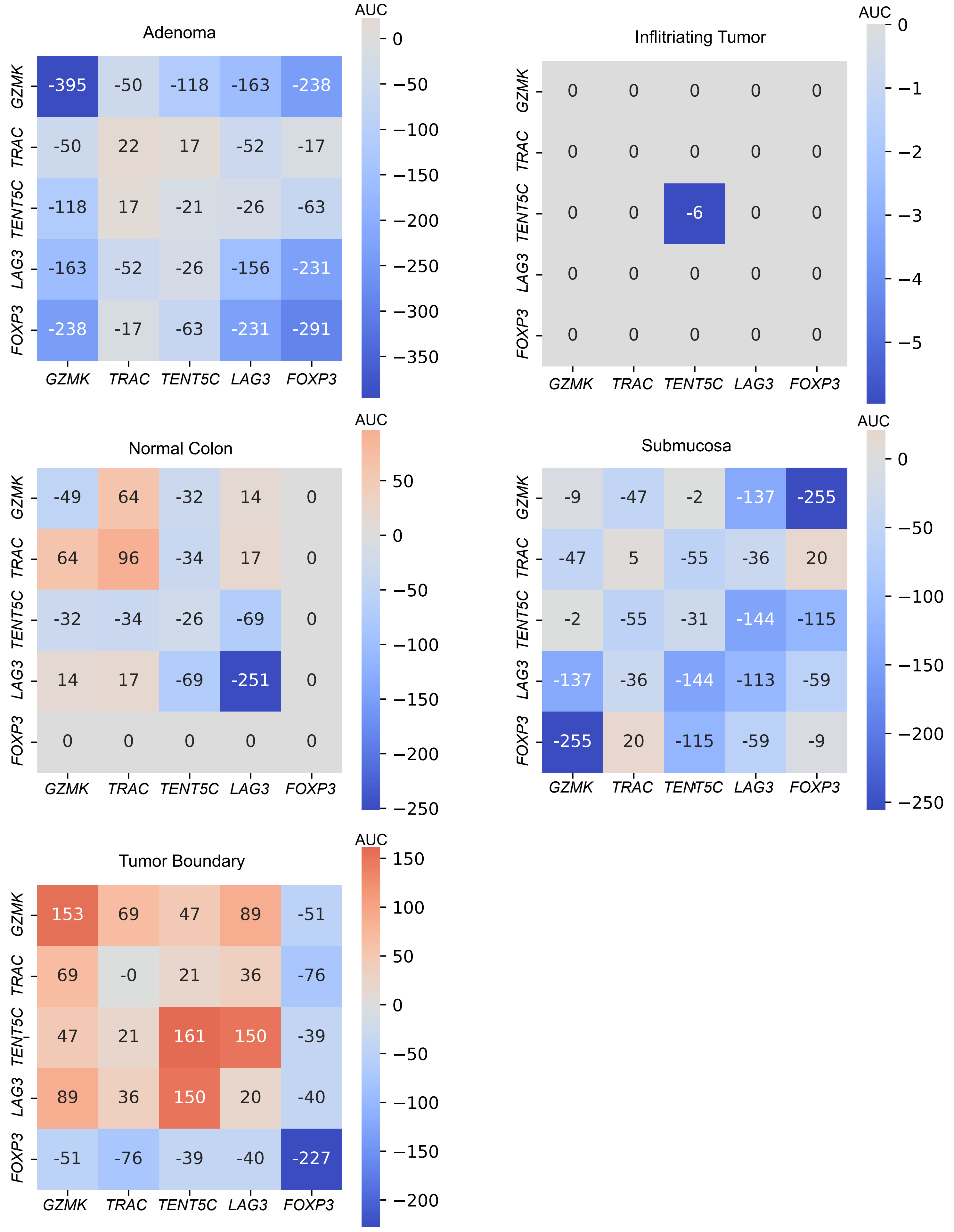
Adenoma
AUC
0
−50
−100
−150
−200
−250
−300
−350
GZMK
TRAC
TENT5C
LAG3
FOXP3
-395 -50 -118 -163 -238
-50 22 17 -52 -17
-118 17 -21 -26 -63
-163 -52 -26 -156 -231
-238 -17 -63 -231 -291
GZMK
TRAC
TENT5C
LAG3
FOXP3
Inflitriating Tumor
AUC
0
−1
−2
−3
−4
−5
GZMK
TRAC
TENT5C
LAG3
FOXP3
0 0 0 0 0
0 0 0 0 0
0 0 -6 0 0
0 0 0 0 0
0 0 0 0 0
GZMK
TRAC
TENT5C
LAG3
FOXP3
Normal Colon
AUC
50
0
−50
−100
−150
−200
−250
GZMK
TRAC
TENT5C
LAG3
FOXP3
-49 64 -32 14 0
64 96 -34 17 0
-32 -34 -26 -69 0
14 17 -69 -251 0
0 0 0 0 0
GZMK
TRAC
TENT5C
LAG3
FOXP3
Submucosa
AUC
0
−50
−100
−150
−200
−250
GZMK
TRAC
TENT5C
LAG3
FOXP3
-9 -47 -2 -137 -255
-47 5 -55 -36 20
-2 -55 -31 -144 -115
-137 -36 -144 -113 -59
-255 20 -115 -59 -9
GZMK
TRAC
TENT5C
LAG3
FOXP3
Tumor Boundary
AUC
150
100
50
0
−50
−100
−150
−200
GZMK
TRAC
TENT5C
LAG3
FOXP3
153 69 47 89 -51
69 -0 21 36 -76
47 21 161 150 -39
89 36 150 20 -40
-51 -76 -39 -40 -227
GZMK
TRAC
TENT5C
LAG3
FOXP3

Supplementary Figure S8: T-cell marker aggregation measurements in a sample of colorectal cancer measured with Visium HD from Figure 5a in different regions of interests. Red: aggregation, blue: repulsion, gray 0 value: no neighboring cells of the specified type.

18

### Supplementary Tables

Supplementary Table S1: Clinical and molecular characteristics of biopsy samples. Abbreviations: MSC, mesenchymal stem cell; NPM1, nucleophosmin 1; FLT3, FMS-like tyrosine kinase 3; ITD, internal tandem duplication; TKD, tyrosine kinase domain; CR1, complete remission 1; ND, not determined; NA, not applicable.

| Biopsy Number | Sex | Age | MSC density | NPM1 | FLT3-ITD | FLT3-TKD | CR1-Achieved | Relapse |
|---|---|---|---|---|---|---|---|---|
| B13-8354 | M | 52 | Low | + | - | - | Yes | No |
| B13-8907 | F | 42 | Normal | ND | ND | ND | Yes | No |
| B13-12681 | F | 53 | Low | ND | ND | ND | NA | NA |
| B14-2846 | M | 77 | Normal | - | + | - | Yes | Yes |
| B14-10782 | M | 68 | Normal | ND | ND | ND | Yes | Yes |
| B14-13014 | F | 29 | Low | + | Intermediate | - | Yes | No |
| B10-2931 | M | 70 | Normal | + | - | - | Yes | Yes |
| B10-13170 | M | 73 | Normal | - | - | - | NA | NA |
| B10-9552 | F | 76 | High | + | - | - | NA | Yes |
| B17-0644 | F | 21 | Normal | ND | ND | ND | No | No |
| B17-6250 | M | 26 | Low | ND | ND | ND | Yes | Yes |
| B17-5404 | M | 28 | Low | ND | ND | ND | Yes | No |

Supplementary Table S2: Normal bone marrow samples.

| Sample code | Age | Sex |
|---|---|---|
| 06-14 | 61 | M |
| 17-16 | 60 | F |
| 11-16 | 65 | F |
| 15-16 | 70 | F |
| 22-16 | 92 | F |

Supplementary Table S3: IMC antibody summary.

| Metal | Antibody | Species | Clone | Cat. Number | Brand | Dilution |
|---|---|---|---|---|---|---|
| 141Pr | alpha SMA | Mouse | 1A4 | 3141017D | Fluidigm | 1 to 5000 |
| 163Dy | CD163 | Rabbit | Polyclonal | ab199402 | Abcam | 1 to 10 |
| 161Dy | CD20-161Er | Mouse | H1 | 3161029D | FLuidigm | 1 to 150 |
| 170Er | CD3 | Rabbit | Poly Rabbit C-term | 3170019D | Fluidigm | 1 to 150 |
| 145Nd | CD31 | Mouse | C31.3 + C31.7 + C31.10 | LS-C390863 | LSBio | 1 to 250 |
| 158Gd | CD34 | Mouse | QBEND/10 | NB120-963 | Novous | 1 to 100 |
| 152Sm | CD45h | Mouse | 2B11 | 3152016D | Fludigm | 1 to 250 |
| 169Tm | Collagen type 1 | Goat | Polyclonal goat | 3169023D | Fludigm | 1 to 5000 |
| 168Er | Ki67 | Mouse | B56 | 3168022D | Fluidigm | 1 to 50 |
| 146Nd | MPO | Rabbit | E1E7I | 14569BF | CST | 1 to 2000 |
| 143Nd | Vimentin | Rabbit | D21H3 | 3143027D | Fluidigm | 1 to 100 |
| 149Sm | CD38 | Rabbit | Polyclonal | HPa022132 | Sigma | 1 to 1000 |
| 142Nd | Perilipin | Rabbit | Polyclonal | ab3526 | Abcam | 1 to 100 |
| 153Eu | WNT3A | mouse | 3A6 | ab81614 | Abcam | 1 to 100 |
| 160Gd | NGFR | Mouse | MLR2 | MA1-18401 | Invitrogene | 1 to 50 |
| 167Er | CXCL-12 | Mouse | 79018 | MAB350 | R&D Systems | 1 to 10 |
| 176Yb | CD10 | Mouse | 56C6 | ab951 | Abcam | 1 to 200 |
| 151Eu | Osteocalcin | Mouse | # 190125 | MAB1419-SP | R&D Systems | 1 to 800 |
| 144Nd | CD90h | Rabbit | EPR3132 | ab221607 | Abcam | 1 to 100 |
| 166Er | c-kit h and m | Goat | polyclonal | AF1356 | R&D SYSTEMS | 1 to 400 |
| 159Tb | CD68 | Mouse | KP1 | 3159035D | Fluidigm | 1 to 2500 |
| 174Yb | E-Cadherin | Rabbit | 24E10 | 3195 | Cell Signaling Technology | 1 to 200 |

Supplementary Table S4: Cox proportional hazards regression analysis results after feature selection based on ranking of individual covariate significance. The exponentiated coefficients correspond to hazard ratios. Statistically significant $p < 0.05$ values are in bold.

| Covariate | Coef | Exp(Coef) | *p* value |
|---|---:|---:|---:|
| **CD34 vs. MPO** | **0.0046251** | **1.0046358** | **0.0328** |
| CD68 vs. CD68 | -0.0065144 | 0.9935068 | 0.7051 |
| CD68 vs. CD31 | -0.0330728 | 0.9674681 | 0.1384 |
| NGFR vs. CD163 | -0.0163394 | 0.9837934 | 0.3205 |
| CD163 vs. Ki67 | 0.0285480 | 1.0289594 | 0.0901 |
| Age | 0.0535937 | 1.0550559 | 0.1164 |
